# Long-read cross-platform validation reveals novel repeat features in myotonic dystrophy type 2

**DOI:** 10.64898/2026.06.06.730578

**Authors:** Marco Carlomagno, Francisco Javier Suárez-López, Simone Maestri, Alfonso Esposito, Vanja Obadović, Virginia Veronica Visconti, Dario Ciabini, Luca Marcolungo, Niccolò Rossi, Mauro Casagrande, Luca Angheben, Lorenzo Spadoni, Maria Rosaria D’Apice, Giuseppe Novelli, Massimo Delledonne, Annalisa Botta, Marzia Rossato

## Abstract

The broader application of long-read sequencing (LRS) for repeat expansion characterization in myotonic dystrophy type 2 (DM2) and other repeat expansion disorders (REDs) remains limited by the lack of systematic validation and benchmarking of sequencing results and bioinformatic workflows.

Here, we performed an orthogonal cross-platform validation of previously generated Oxford Nanopore Technologies (ONT) data by sequencing the same DNA samples with Pacific Biosciences (PacBio) HiFi following amplification-free targeted enrichment in a cohort of 8 DM2 patients. Despite substantial differences in sequencing chemistry and coverage, the two platforms showed high concordance in repeat size estimation, somatic mosaicism, and repeat architecture. This validation confirmed the presence of the (TCTG)n motif and enabled the identification of a previously unreported (CCCG)n motif at the 3′ end of expanded alleles, further highlighting the structural complexity of the *CNBP* expansion. Through this analysis, we also established a bioinformatic workflow that improved ONT-based repeat characterization, addressing limitations in motif resolution and enabling more accurate analysis of *CNBP* expansions.

Overall, this study provides a validated framework for LRS-based *CNBP* repeat analysis, supporting the integration of these technologies into routine molecular investigation for DM2 and other REDs.

## INTRODUCTION

Over the past decade, long-read sequencing (LRS) has fundamentally transformed the investigation of repeat expansion disorders (REDs). By enabling direct sequencing across long, GC-rich, and highly repetitive regions, LRS overcomes many of the limitations of amplification-based approaches traditionally used for RED molecular diagnosis 1,2. Beyond improving repeat sizing accuracy, LRS has revealed previously inaccessible structural features, including novel repeat motifs, sequence interruptions, and complex motif architectures3–5. In several REDs, such features have proven biologically relevant, refining disease classification and uncovering modifiers of disease onset and progression6,7. However, the routine application of these technologies for RED characterization and diagnosis is currently limited by the lack of systematic validation and benchmarking of LRS results, bioinformatic workflows, and data interpretation8 (Benarroch et al, Nature Genetics 2026, in revision; PMID: 39676657).

Myotonic dystrophy type 2 (DM2; MIM#602668) is an autosomal dominant multisystemic disorder exemplifying both the challenges and the impact of this technological shift. DM2 is caused by a large and unstable (CCTG)n expansion within intron 1 of the *CNBP* gene, typically arranged in a complex (TG)*v*(TCTG)*w*(CCTG)*n* configuration^9,10^. These expansions can reach up to ∼40 kb, and exhibit pronounced somatic mosaicism, rendering conventional PCR-based methods unreliable and Southern blot only partially informative8,11,12. In previous work, we applied amplification-free Cas9-mediated enrichment combined with Oxford Nanopore Technologies (ONT) sequencing to characterize *CNBP* repeat expansions at single-molecule resolution13,14. This approach enabled full-length sequencing of expanded alleles, quantitative assessment of somatic mosaicism, and the identification of a previously unreported (TCTG)n motif embedded within the expanded tract in most of DM2 patients13. These findings expanded the structural landscape of DM2 and highlighted the potential of LRS to improve the precision of *CNBP* repeat characterization in a clinical context. However, robust validation of the identified repeat features through orthogonal approaches remains essential, given the potential impact of these findings on disease prognosis.

The presence of the (TCTG)n motif in *CNBP* expansions has been confirmed in independent studies using long-read sequencing and by QP-PCR analyses in larger DM2 cohorts 14,15. Nevertheless, substantial uncertainty remains regarding its precise sequence features, including motif length and the presence of interruptions. To date, only a small number of DM2 patients (20) have been analyzed by LRS, with most data generated by our research group13,14, and an independent dataset characterized by limited coverage of the expanded allele in a single patient15,16. In addition, ONT-based analyses frequently yielded sequence segments without clear repeat motif annotations within expanded alleles -particularly within the (TCTG)n tract-thereby limiting the ability to confidently define motif composition14. This ambiguity raises the possibility that, beyond (TCTG)n and (CCTG)n motifs, additional repeat sequences or interruptions may be present within the expanded tract. Since sequencing of full-length expanded alleles in DM2 has so far relied exclusively on ONT technology, at least part of this uncertainty may reflect technical limitations inherent to ONT sequencing and associated bioinformatic analyses. While ONT offers a major advantage in generating ultra-long reads without intrinsic length limits -making it particularly suitable for large expansions like those in *CNBP*-it is also characterized by higher error rates compared to other sequencing technologies. These errors, particularly insertions and deletions, are enriched in homopolymeric and repetitive regions17–19. Although recent improvements in sequencing chemistry and basecalling algorithms have significantly increased raw-read accuracy, error rates remain non-negligible in repeat-dense regions16,20. The presence of systematic errors can hinder the accurate detection of specific motifs, such as (TCTG)n, and may lead to the misidentification of novel motifs, while also impacting the reliable quantification of repeat mosaicism. For very long molecules, even modest per-base error rates can accumulate, potentially distorting repeat-length estimates and leading to under-or overestimation.

In the absence of orthogonal validation, it remains thus unclear to what extent newly described structural features reflect true biological variation versus ONT-specific sequencing or even basecalling artifacts. Previous studies in another RED, myotonic dystrophy type 1 (DM1, MIM#160900), have shown that different basecalling algorithms can yield divergent results, underscoring the importance of carefully evaluating bioinformatics pipelines for ONT-based data analysis1,16,21. In particular, comparative analyses showed that different basecallers, including the ONT production basecaller Dorado and the trainable neural network basecaller Bonito, yield substantially different results in repeat size estimation, sequence composition, and mosaicism profiles16. These observations suggest that alternative or optimized basecalling strategies may improve repeat characterization in DM2.

More broadly, standardized workflows to fully exploit the potential of LRS in repeat expansion analysis remain lacking, both for DM2 and for other REDs. The implementation of robust pipelines in research and clinical settings, therefore, requires systematic orthogonal validation of LRS results, along with comprehensive benchmarking of both experimental approaches and bioinformatic pipelines. Together with ONT, Pacific Biosciences (PacBio) single-molecule real-time sequencing is the leading LRS technology capable of spanning large repeat expansions22,23. PacBio exhibits a distinct error profile compared to ONT, with high per-base accuracy (Q>30) achieved through circular consensus (HiFi) sequencing, albeit at generally shorter read lengths (∼20 kb on average) 24,25. These complementary features make ONT and PacBio ideally suited for cross-platform validation of complex repeat structures. However, PacBio has not yet been systematically applied to DM2 repeat expansions, and direct cross-platform comparisons remain lacking.

In this study, we performed an orthogonal cross-platform validation of previously generated Oxford Nanopore Technologies (ONT) data using Pacific Biosciences (PacBio) HiFi sequencing in a cohort of eight DM2 patients. This comparison is particularly timely given the recent release of the PacBio PureTarget kit, which enables targeted analysis of multiple repeat loci, including *CNBP*, using a Cas9-mediated enrichment. By analysing matched datasets using a consistent bioinformatic framework, we aimed to assess the robustness of previously reported DM2 repeat features and to directly evaluate the performance of the two platforms in resolving repeat size, somatic mosaicism, and internal repeat architecture. This approach also enabled the identification of previously unrecognized repeat features and the optimization of analytical strategies for the characterization of complex repeat expansions.

## RESULTS

### *CNBP* repeat-expansion sequencing using Cas9-mediated enrichment coupled to PacBio sequencing

To cross-validate long-read sequencing analysis of *CNBP* repeat expansions in DM2, we selected eight patients from our previous study, for whom ONT data had already been generated using Cas9-mediated enrichment14, including two with pure (CCTG)n expansions and six harburing the (TCTG)n motif. The same DNA samples were sequenced using PacBio HiFi with the PureTarget kit. For both ONT and PacBio sequencing, Cas9-enrichment of the CNBP repeat locus used identical gRNAs targeting a 4.2 kb region of the human reference genome. PacBio sequencing generated a median of 65,459 HiFi reads per sample, of which 2,597 mapped to the CNBP locus (median on-target rate: 3.58%; **Supplementary Table 1**), resulting in a median coverage of 320x for the expanded allele, compared with 84x previously obtained with ONT14.

Both datasets consistently showed an underrepresentation of reads derived from the expanded allele across all patients, with a median of 6.9 times lower read counts compared to the normal allele (median fraction of 13.13% for ONT and 14.55% for PacBio; **Figure 1A**). The fraction of reads derived from the expanded allele was inversely proportional to its median expansion length, suggesting a lower sequencing efficiency for longer alleles in both technologies (**Figure 1B**), as already observed in DM2 and other REDs26,27. Consistently, no allele-specific bias occurred at the not-expanded control locus *FMR1*, present in both gRNA panels, where reads were distributed evenly between the two alleles on both platforms (**Supplementary Figure 1**).

**Figure 1.**
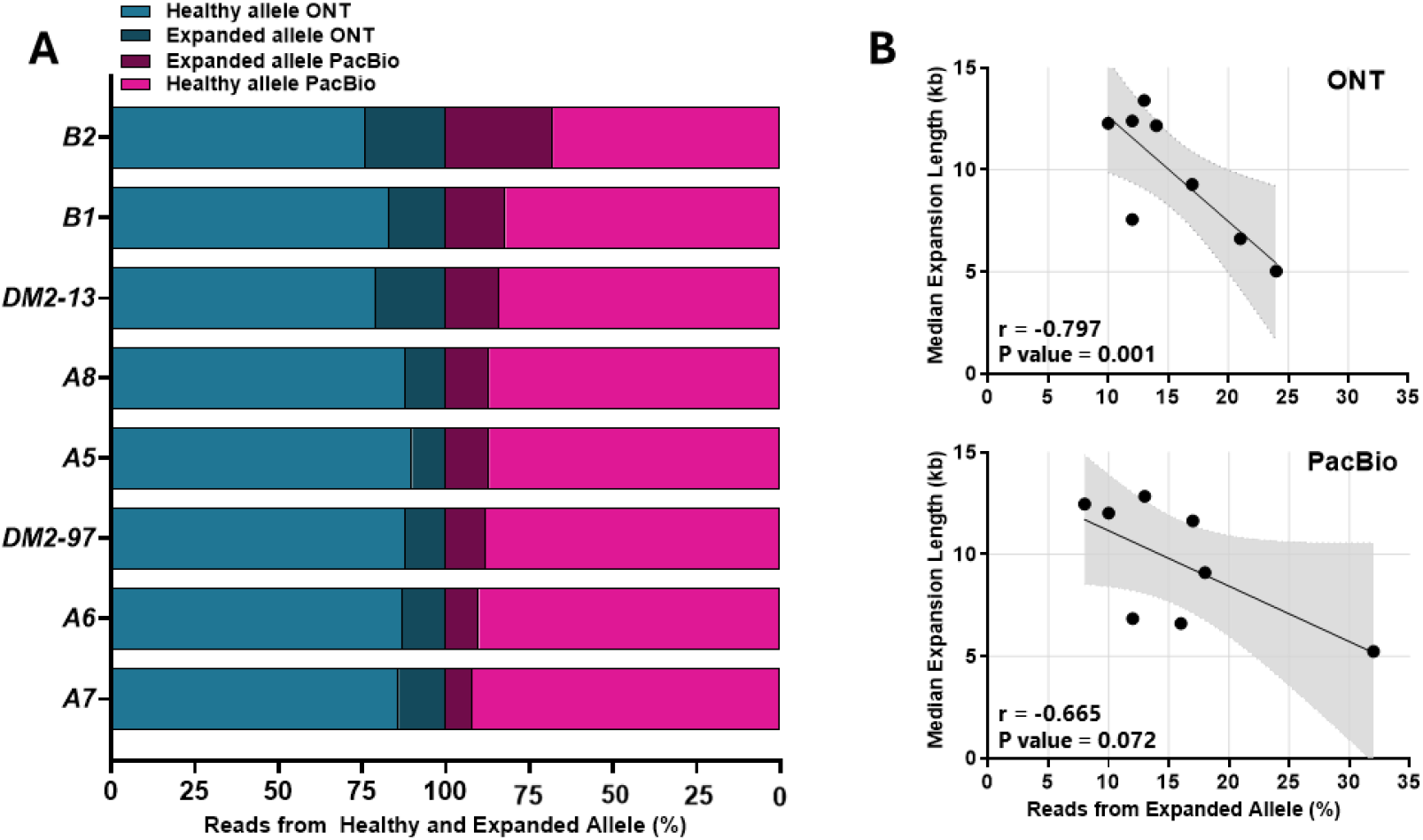
Sequencing data from *CNBP* normal and expanded alleles in DM2 patients. **A**) Bar plots showing the proportion of reads mapping on *CNBP* and assigned to the normal and expanded alleles for each sample using ONT and PacBio. Numbers in each bar fraction indicate the absolute number of reads assigned to each allele. Patients are ranked based on the median size of expanded alleles. **B**) Scatter plots showing the negative correlation between the fraction of reads from the expanded allele and the median expansion length measured based on ONT and PacBio sequencing data for each patient. The black line indicates the linear regression fit, and the light grey shadow its 95% confidence interval. Pearson correlation was used to assess the linear association between the two variables.

### *CNBP* repeat-expansion size analysis

PacBio data revealed substantial variability in the size of reads originating from expanded alleles across all patients (**Figure 2A** and **Figure 3A**), confirming the high degree of somatic mosaicism previously observed with ONT sequencing14. Expansion lengths ranged from 0.43 to 32.07 kbp, corresponding to up to 107-8,017 repeat units, and were similar to those previously reported using ONT (0.45 to 40.29 kbp14). Despite differences in sequencing coverage, Kolmogorov–Smirnov tests confirmed no statistically significant differences in repeat size distributions between the two platforms accross all patients, except for patient A7 (**Figure 2A**).

**Figure 2.**
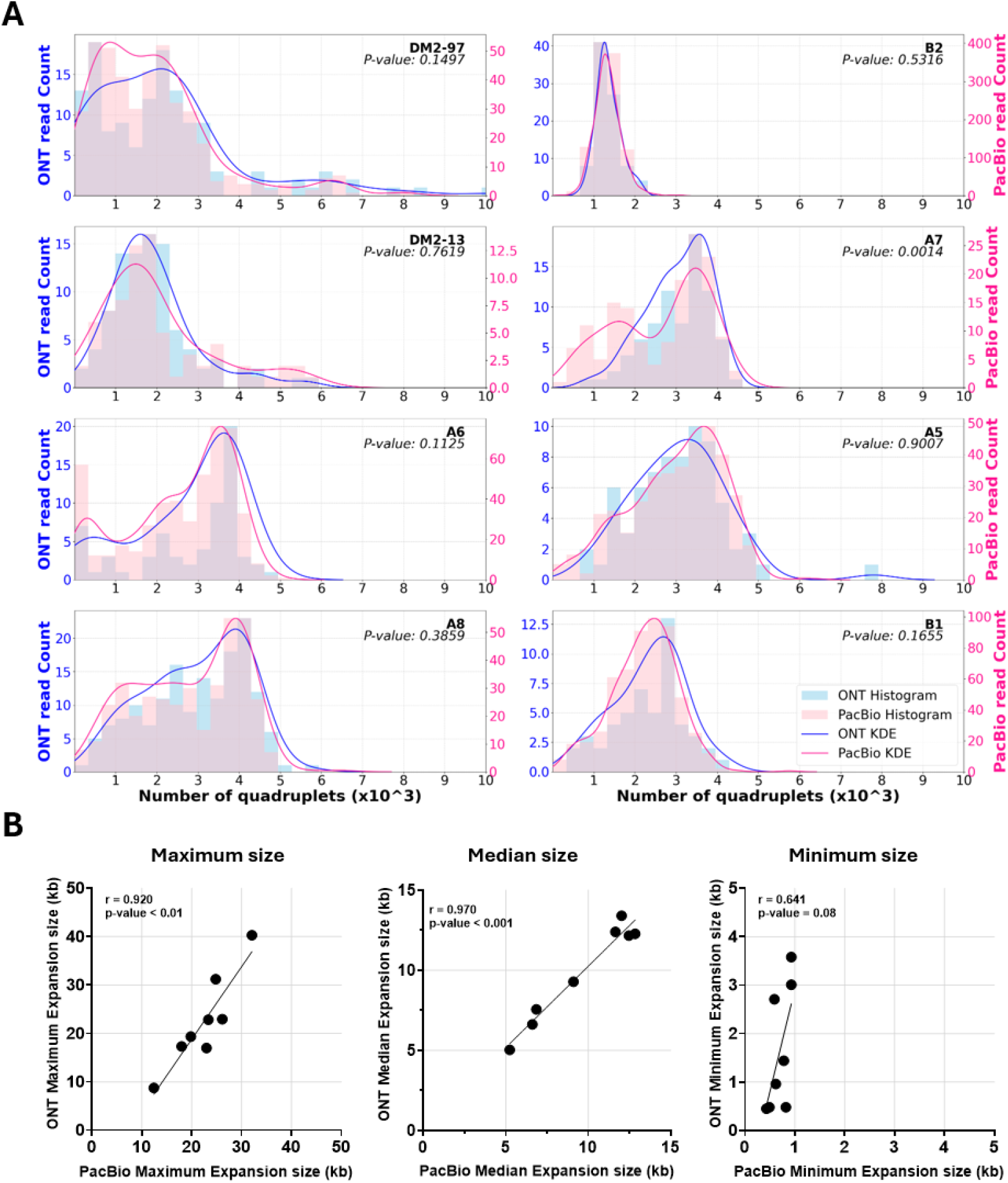
Size characterization of expanded *CNBP*-repeat alleles in DM2 patients. **A**) Abundance distribution of quadruplets identified in each DM2 patient. The y-axis shows the number of ONT/PacBio reads with a certain number of repeats, whereas the x-axis shows the number of quadruplet repeats identified. Reads were grouped into 1320 bp bins (330 quadruplets). Lines represent the estimated kernel density of the underlying solid distribution of ONT/PacBio reads. P-values indicate the results of the Kolmogorov–Smirnov tests comparing the two distributions. **B**) Correlations between the maximum, median and minimum expansion lengths detected by ONT and PacBio sequencing for each DM2 patient. Pearson correlation was used to assess the linear association between the two datasets.

**Figure 3.**
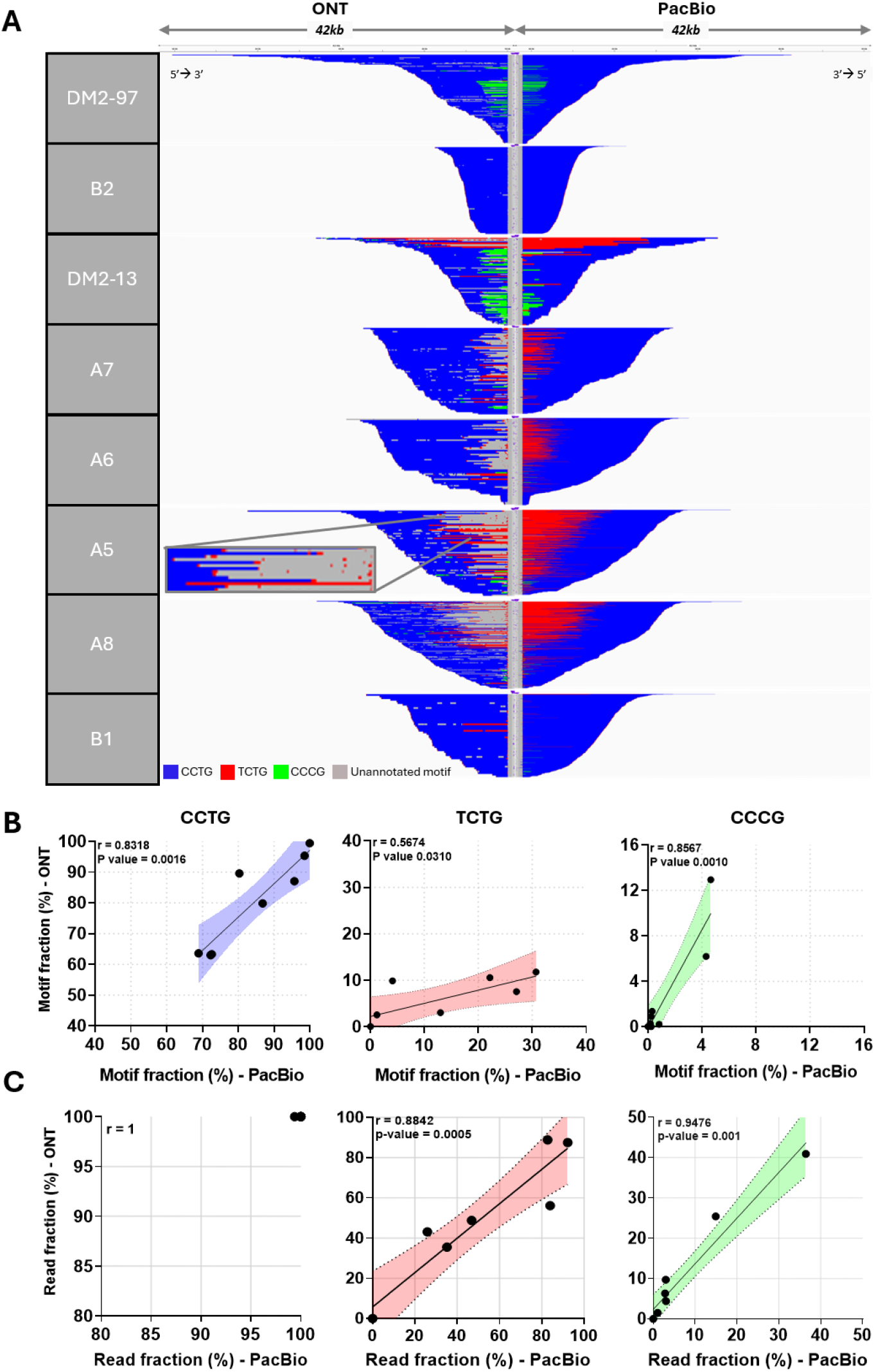
Sequence structure of expanded *CNBP* alleles in DM2 patients. **A**) Integrative Genomics Viewer (IGV) visualization (42-kbp window) of ONT (left) and PacBio (right) targeted sequencing data from *CNBP* expanded alleles of each DM2 patient. Complete reads were aligned at the 3′ end of the repeat in order to identify the repeat pattern that characterizes the expanded microsatellite locus. Each motif in the expanded alleles was visualized using a different colour, as indicated in the key. The inset shows a zoomed-in view of the indicated read segment, highlighting the lower resolution of ONT in detecting the (TCTG)n motif. **B**) Correlations between the fraction of repeat units across the total number of repeats in ONT and PacBio datasets, displaying the CCTG motif (left panel), the TCTG motif (central panel) and the CCCG motif (right panel), for each DM2 patient. **C**) Correlations between the fraction of ONT and PacBio reads displaying the CCTG motif (left panel), the TCTG motif (central panel) and the CCCG motif (right panel), for each DM2 patient. Pearson correlation was used to assess the linear association between the two datasets.

To assess whether the observed variability in molecule size reflected genuine biological heterogeneity or technical artifacts, we extracted PacBio subreads from representative reads of different lengths from patient DM2-13. Subread length measurements closely matched those of the corresponding HiFi consensus sequences, showing near-perfect concordance (**Supplementary Figure 2**) and indicating that the observed variability is not introduced during the consensus-calling process used to generate HiFi reads.

Since accurate estimation of expansion size could be relevant for genotype-phenotype correlation studies and disease prognosis, we compared the maximum, median and minimum *CNBP* repeat lengths detected by ONT and PacBio. Maximum and median length showed excellent cross-platform concordance, with correlation coefficients close to 1 (r = 0.920 and r= 0.970, respectively), both highly significant (p < 0.01) (**Figure 2B**). In contrast, minimum expansion lengths showed only moderate correlation (r = 0.641, p = 0.08) and were longer in ONT data for 5 patients, suggesting that estimation of the smallest expanded molecules may be influenced by platform-specific factors (**Figure 2B**). To assess whether this difference could be attributable to the lower coverage in the ONT dataset, we downsampled PacBio reads to match the number of ONT reads for each expanded allele and re-estimated the expansion length. After down-sampling, the difference between PacBio and ONT minimum repeat lengths decreased from 2.21-to 1.33-fold on average across all patients (**Supplementary Figure 3**); however, the correlation remained non-significant, suggesting that reduced ONT coverage only partially explains the overestimation of minimum expansion length.

### *CNBP* repeat motif and structure analysis

To assess the sequence composition and structure of *CNBP* repeat expansions, previously generated ONT reads from expanded alleles14 were re-analysed alongside the PacBio dataset using a unified bioinformatic pipeline based on MosaicViewer, as described in the method section. The two sequencing platforms showed qualitatively concordant results in the repeat structure (**Figure 3A**). Specifically, all patients presented the expected (TG)v(TCTG)w(CCTG)n structure at the 5’ end of the repeat locus. However, only one patient (B2) featured a ‘pure’ pattern of (CCTG)n repeats until the 3’ end of the repeat. Consistent with previous reports13–15, the majority of patients (6 out of 8) presented long stretches of (TCTG)n arrays (coloured in red) at the 3’ end of the CCTG expansion (**Figure 3A**). The TCTG motif was more robustly detected in PacBio reads, whereas ONT datasets showed a higher proportion of read segments with unannotated repeat motifs in the corresponding regions (coloured in gray, **Figure 3A**), indicating an intrinsic technical limitation of ONT in resolving this motif, as previously observed14. Interestingly, we also consistently identified another atypical motif, (CCCG)n (coloured in green), at the 3’ end of the expansion with both technologies (**Figure 3A**). To our knowledge, this CCCG repetition was never reported in *CNBP* expansions before. The (CCCG)n motif was detected by both platforms in all (TCTG)n positive patients, showing the co-existence of (TG)*v*(TCTG)*w*(CCTG)*n*(TCTG)*m* and (TG)*v*(TCTG)*w*(CCTG)*n*(CCCG)*m* repeated arrays in different reads of the same patient (**Figure 3A**). Finally, a single patient (DM2-97) presented only the atypical (CCCG)n motif at the 3’ end, in addition to the expected (CCTG)n stretch, with no additional (TCTG)n sequences (**Figure 3A**).

To compare the sensitivity of the two platforms in detecting the three motifs characterizing the *CNBP* expansions, we evaluated both their overall abundance -defined as the fraction of repeat units across the total number of repeats- and their frequency at the read level, i.e., the fraction of reads carrying each motif (**Figure 3B** and **3C**). Overall, both platforms showed consistent detection of the three motifs, as indicated by strong correlations in both motif abundance and read-level fraction. When present, the (TCTG)n and the (CCCG)n motifs were detected in a highly variable fraction of sequences across donors, and differed widely in length both within and across donors (**Figure 3A** and **Supplementary Figure 4**). However, compared to PacBio, ONT tended to underestimate the abundance of (TCTG)n motifs (**Figure 3A** and **3B**).

**Figure 4.**
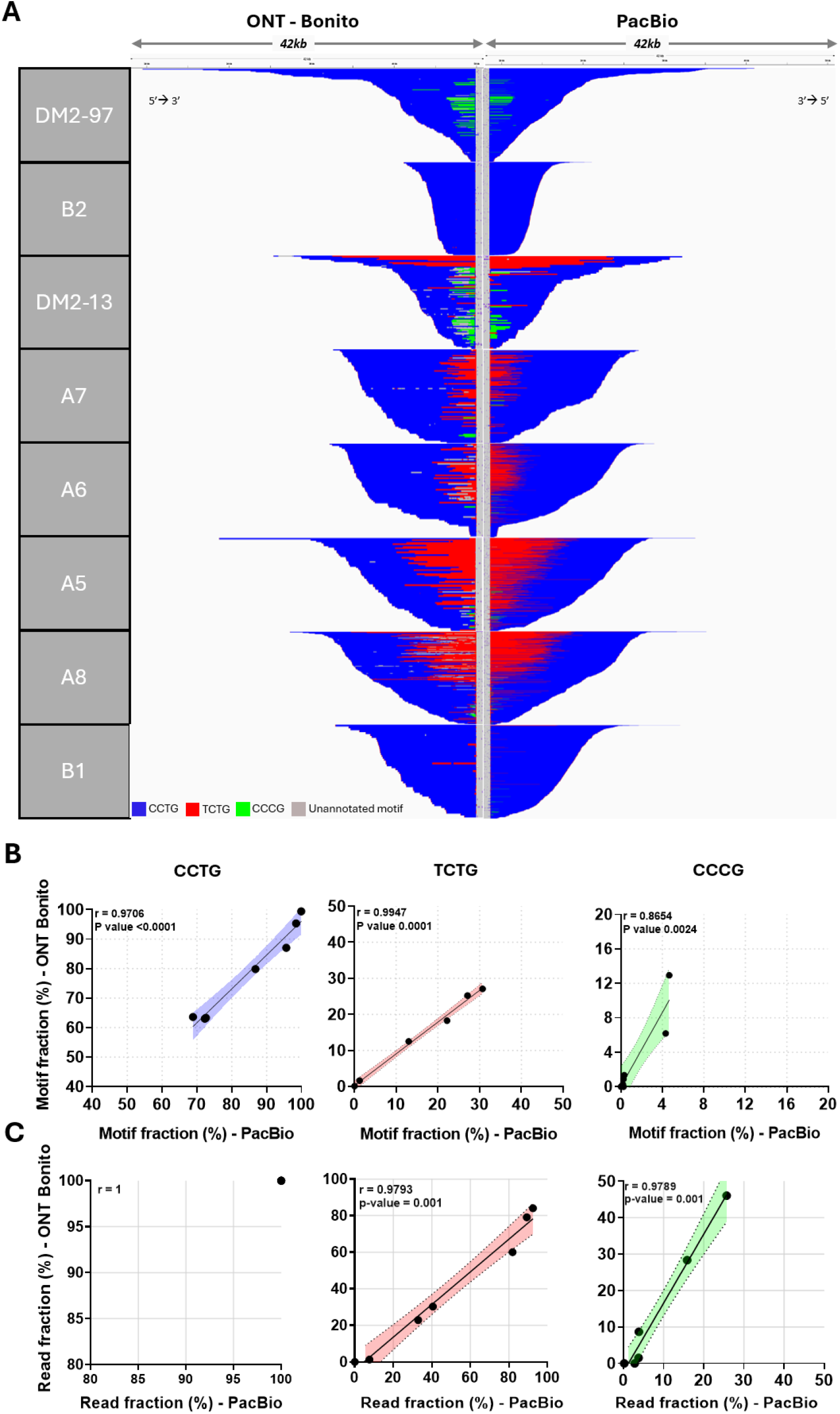
Sequence structure of expanded *CNBP* alleles based on ONT data re-basecalled with Bonito and PacBio data. Comparison of PacBio data with ONT data re-basecalled using Bonito trained on PacBio data with the Leave-One-Out (LOO) strategy. **A**) Integrative Genomics Viewer (IGV) visualization (42-kbp windows) of expanded *CNBP* alleles from ONT-Bonito LOO (left panel) or PacBio sequencing data (right panel). Complete reads were aligned at the 3′ end of the repeat, in order to identify the repeat pattern that characterizes the expanded repeat locus. Each motif in the expanded alleles was visualized using a different colour, as indicated in the key. **B**) Correlations between the fraction of repeat units across the total number of repeats in the ONT and PacBio data, displaying the CCTG motif (left panel), the TCTG motif (central panel) and the CCCG motif (right panel), for each DM2 patient. **C**) Correlations between the fraction of ONT and PacBio reads displaying the CCTG motif (left panel), the TCTG motif (central panel) and the CCCG motif (right panel), for each DM2 patient. Pearson correlation was used to assess the linear association between the two datasets. The same PacBio data are also shown in Figure 3.

### Refinement of *CNBP* repeat motif analysis based on ONT sequencing

The limited ability of ONT data to accurately resolve the (TCTG)n motif represents a potential constraint for its application in *CNBP* repeat analysis, particularly in a diagnostic context, as it introduces uncertainty in motif assignment. To assess whether this effect may be attributable to basecalling-related inaccuracies, we re-basecalled ONT raw data using Bonito, a trainable basecaller that can be optimized using external datasets. Specifically, the Dorado model previously used for ONT basecalling was fine-tuned within the Bonito framework using PacBio data from all patients except one, and the resulting models were applied to re-basecall ONT data from the held-out sample (Leave-One-Out (LOO) strategy). Bonito-based fine-tuned basecalling improved motif resolution compared to standard ONT basecalling with Dorado or non-fine-tuned Bonito, as highlighted by a consistent reduction in the fraction of bases without motif assignment (**Supplementary Figure 5** and **6**). Consistently, quantitative agreement between PacBio and ONT data improved in both repeat-level abundance and read-level frequency compared to pre–re-basecalling analyses (**Figure 4B** and **C**). Importantly, the re-basecalling approach did not introduce detectable artifacts, as no additional motifs were observed in patients who originally presented with only one or two motif types (**Figure 4; Supplementary Figures 5** and **6**). Moreover, they did not alter basecalling accuracy outside the *CNBP* repeat region, as no increase in error rates was observed in the other two loci included in the Cas9 enrichment panel (**Supplementary Table 2**). Overall, these results suggest that basecalling plays a key role in accurately identifying repeat motifs with ONT sequencing and indicate that, when appropriately processed, ONT data can achieve accuracy comparable to PacBio for characterizing *CNBP* repeat architecture.

## DISCUSSION

Here, we provide orthogonal validation of ONT data using PacBio HiFi sequencing to comprehensively characterize *CNBP* repeat expansions in DM2. Despite fundamental differences in sequencing chemistry and on-target coverage achieved, the two platforms converged on concordant estimates of expansion size, somatic mosaicism, and repeat architecture, including the identification of a novel motif at the 3’ end of the (CCTG)n array.

With the experimental setup used (8-plex on a Revio SMRT Cell and up to three samples per MinION flow cell), both platforms consistently achieved >50x coverage of the expanded allele across all individuals, exceeding the minimal suggested threshold for clinical repeat expansion analysis28. However, PacBio achieved a higher on-target fraction and generated approximately threefold more CNBP-mapping reads per sample. Higher ONT coverage could be achieved by extending sequencing time, increasing DNA input, or pooling multiple libraries from the same sample within a single sequencing run. Importantly, only a small fraction of available pores or SMRT wells was occupied on both platforms, indicating that sequencing output is primarily limited by the number of adapted molecules rather than by flow cell capacity. This suggests that improved Cas9-enrichment efficiency and optimized multiplexing strategies could substantially increase throughput and reduce costs. Current ONT Cas9-based multiplexing approaches, however, remain relatively inefficient13,15. Higher multiplexing levels are technically feasible with PacBio but require reduced DNA input per sample. Given the pronounced allelic imbalance observed, this trade-off should be carefully evaluated, as lower DNA input may disproportionately reduce coverage of expanded alleles.

A systematic and marked underrepresentation of *CNBP* expanded alleles relative to normal alleles was observed with both technologies, with the fraction of expanded reads inversely correlated with repeat length. This may possibly reflect preferential recovery and sequencing of shorter DNA fragments, as longer molecules are more susceptible to fragmentation during sample preparation and intrinsically more difficult to capture and sequence, even with amplification-free approaches. Similar biases have been reported in our previous work on DM213 and in other genes associated with REDs, including *DMPK*29 and *C9orf72*30,31, despite the generally shorter length of expanded alleles at these loci. Notably, this bias was observed to a similar extent in both ONT and PacBio despite their fundamentally different sequencing chemistries, raising the possibility that additional factors may contribute to this phenomenon beyond technical limitations in the enrichment or sequencing of very large or structurally complex DNA molecules.

While median and maximum expansion size estimates were highly correlated between platforms, ONT tended to overestimate the minimum size of expanded alleles. This effect was likely attributable to lower sequencing coverage and could potentially be mitigated by increasing data yield, as described above. Accurate estimation of the minimum expansion size is critical for understanding somatic mutational dynamics and defining the progenitor allele length (ePAL), a clinically relevant parameter, as exemplified in DM1 where it is a major determinant of disease severity32. Currently, assessment of ePAL relies on technically demanding methods such as small-pool PCR (SP-PCR)33, which are not readily scalable for routine diagnostics in DM2. Given that LRS represents a promising alternative, the ability of these sequencing technologies to accurately estimate ePAL warrants further investigation in dedicated studies.

Both platforms confirmed extreme somatic mosaicism in *CNBP* expansions, as previously observed13. From a practical perspective, these findings highlight the importance of achieving high locus-specific coverage when analyzing large and mosaic expansions to capture full-size variability. From both a biological and clinical perspective, it remains important to determine whether comparable levels of somatic instability are present across disease-relevant tissues, including skeletal muscle, cardiac and brain tissues, and the central nervous system as described for other REDs, including Huntington’s disease34,35. In these tissues, the distribution of CCTG expansion sizes may differ substantially from that observed in peripheral blood, potentially influencing the phenotypic variability of clinical symptoms in DM2 patients.

This cross-validation enabled robust confirmation of previously reported (TCTG)n motif and, importantly, the consistent identification of a novel (CCCG)n motif at the 3′ end of *CNBP* expanded alleles. Detection across independent sequencing chemistries strongly supports the biological authenticity of this motif rather than a platform-specific artifact. Notably, the (CCCG)n motif was observed in both familial and sporadic cases, suggesting that it may be widespread in *CNBP* expansions, although this requires validation in larger cohorts. This novel finding further highlights the value of LRS for the molecular characterization of the DM2 locus and for resolving the full complexity of repeat expansions. The origin of this motif and its potential impact on disease mechanisms, inheritance, and clinical variability remain to be investigated in larger cohorts and through dedicated functional studies.

Beyond structural and sequence characterization, this validation study also enabled the development of an optimized bioinformatic strategy for ONT-based analysis of *CNBP* repeats, exploiting the trainable Bonito basecaller. While a similar strategy has been explored in other genomic contexts, including plant genomes36 and highly repetitive regions such as centromeres and telomeres37,38, its application to pathogenic repeat expansions had not been previously demonstrated. These results thus establish a proof of principle that re-tuned, PacBio data-driven basecalling strategies can be systematically applied to improve ONT-based analysis of pathogenic repeat expansions, providing a generalizable framework for other challenging REDs. Although all re-tuned Bonito models improved detection of the (TCTG)n motif, their performance varied across training datasets, but the limited size of the current DM2 cohort precluded identification of the factors underlying these differences. Extending this approach to larger cohorts will be essential to systematically identify the determinants of model performance, and the present findings should therefore be interpreted as preliminary evidence supporting this strategy. In the future, aggregated high-quality datasets could be used to develop disease-specific basecalling models suitable for routine and clinical applications based on ONT sequencing.

Taken together, our findings suggest that PacBio represents the method of choice for the discovery and high-resolution characterization of novel repeat motifs, owing to its high base accuracy, whereas ONT -once appropriately optimized and validated-provides a flexible solution for routine analysis. From a practical perspective, ONT platforms offer significant advantages in cost and ease of implementation, requiring less infrastructure and a lower initial investment than PacBio systems. For small sample sizes or sequencing individual samples, ONT remains the most cost-effective solution, as the current PacBio PureTarget workflow requires simultaneous processing of at least 8 samples. For larger batches (≥ 8 samples), PacBio becomes more cost- and time-efficient due to higher multiplexing capacity and the availability of a robust, standardized workflow based on a commercial kit and standardized bioinformatic tools. Beyond cross-platform validation, our findings support the immediate clinical applicability of amplification-free long-read sequencing for the molecular diagnosis of DM2, particularly in patients with atypical presentations or negative repeat-primed PCR results. Furthermore, the standardized workflow and orthogonal validation presented here establish a technical foundation that could inform future qualification by regulatory agencies such as the FDA or EMA for use as an in vitro diagnostic device for repeat expansion disorders.

To our knowledge, this is the first systematic cross-platform assessment of amplification-free ONT and PacBio sequencing for analyzing repeat expansions in DM2 and, more broadly, across REDs. By integrating orthogonal validation at both experimental and bioinformatic levels, we demonstrated robust and reliable characterization of *CNBP* repeat expansions, confirming both previously described and newly identified structural features. This validation approach defined the respective strengths and limitations of the two platforms and provided a practical basis for implementing LRS in DM2 diagnosis, while offering a transferable strategy for optimizing repeat-analysis workflows in other REDs.

## MATERIALS AND METHODS

### LRS sequencing

High Molecular Weight (HMW) DNA from DM2 patients was previously extracted14. ONT sequencing data were previously generated14.

Using the same DNA samples, PacBio sequencing data were generated using the PureTarget kit v.1 (PacBio, Menlo Park, CA) according to manufacturer instructions (PN 103-740-700, REV04 JAN2026) and by multiplexing 8 samples in a single library preparation. Briefly, 4ug of HMW DNA was dephosphorylated, and a Cas9-mediated excision was performed to enrich the *CNBP* repeat using the PureTarget Repeat Expansion panel v.1. Excised molecules were ligated to SMRTbell adapter indexes, and an exonuclease treatment was performed to remove off-target molecules. All samples were sequenced on a Revio system using a 25M SMRT Cell with SPRQ reagents, except for sample DM2-13, which was sequenced on a Sequel II. The movie time was set to the default 24 hours using SMRT Link v. 25.2 software.

### Bioinformatic Analysis of *CNBP* Expanded Repeats

ONT POD5 files were basecalled using Dorado (v1.1.1), applying the model of super-high accuracy (SUP) v4.3.0 (dna_r10.4.1_e8.2_400bps_sup@v4.3.0). Downstream analysis was performed only on reads with a minimum quality score of 10. The basecalled reads were aligned to the human reference genome (hg38) using Minimap2 (v2.28) with the -ax map-ont preset, optimized for ONT data. Raw PacBio reads were demultiplexed using SMRT Link v25.2 (Pacific Biosciences, CA, USA) and only HiFi reads were used for downstream analysis. HiFi reads were aligned to the human reference genome (hg38) using pbmm2 v1.17.0 with the HiFi preset. Secondary alignments were excluded for both datasets, retaining only primary mappings.

For both platforms, the resulting alignments were exported in FASTQ format for downstream analyses and analyzed with the same pipeline. Reads fully spanning the *CNBP* repeat region, defined as complete reads, were extracted as previously described13 and analyzed with MosaicViewer (https://github.com/MaestSi/MosaicViewer_CNBP) using the integrated NCRF parameters --minlength 16 and --minmratio 0.75. The novel CCCG motifs was incorporated into the tool to enable their visualization and subsequent analysis.

Simplified reads were generated with MosaicViewer (https://github.com/MaestSi/MosaicViewer_FMR1) as previously described13, with an improved strategy for handling alignment artifacts in the flanking regions. Specifically, a tolerance parameter was introduced to prevent the artificial insertion of blocks of Ns between the expansion start and the flanking sequence. Annotated repeat motifs were visualized in Integrative Genomics Viewer (IGV). The size of the expanded region was estimated by measuring the length of the soft-clipped portion of the CIGAR strings in the BAM files generated by MosaicViewer, and used to generate read-length distributions for the expanded alleles, and to extract maximum, median and minimum values. Motif composition was determined based on motifs identified by NCRF within MosaicViewer. For each sample, motif counts were calculated by extracting the number simplified bases of each type from the MosaicViewer output, and the number of reads displaying each motif was determined by counting the number of reads supporting each motif across the read set.

### Fine-tuning of ONT basecalling

The Dorado basecalling model (dna_r10.4.1_e8.2_400bps_sup@v4.3.0) was fine-tuned within the Bonito framework using PacBio HiFi data. A leave-one-out strategy (LOO) was used, in which, for each patient, the model was fine-tuned based on the data from all other patients and subsequently applied to the excluded individual. An initial basecalling step was performed for each patient using Bonito (v0.8.1) in combination with super-accurate (SUP) v4.3.0 model (dna_r10.4.1_e8.2_400bps_sup@v4.3.0) with the --save-ctc flag enabled, following the recommended training workflow. For each LOO iteration, PacBio HiFi reads were merged using a custom script, according to the Bonito guidelines, excluding the data of patient corresponding to that iteration. Fine-tuning was then performed with Bonito (v0.8.1) using the training datasets generated in the previous step, with default training parameters (learning rate 2e-3, 5 epochs, batch size 64). Finally, ONT POD5 files of each patient were basecalled using the corresponding LOO fine-tuned model trained on all remaining patients, retaining only reads with a minimum quality score of 10. Resulting FASTQ files were analyzed as described in the previous section. Reads corresponding to each target locus were subsequently extracted using Samtools view, and error rates were calculated using Samtools stats.

### Ethical Statement

The analyzed samples were obtained from eight Italian patients of Caucasian ancestry with genetically confirmed DM2 diagnosis. Their participation in the study was approved by the institutional review board of Policlinico Tor Vergata (document no. 232/19). Informed consent was obtained from all patients, and all their clinical information was immediately anonymised. All the experimental procedures were conducted strictly according to the Code of Ethics of the World Medical Association (Declaration of Helsinki).

## Acknowledgement

This work was funded by Ministero della Salute, PNRR: M6/C2_CALL 2022 (GEPINDM project code PNRR-MR1-2022-12375877), and the European Union under the project ENTRY-DM: 101169266-ENTRY-DM-HORIZON-MSCA-2023-DN-01. Views and opinions expressed are, however, those of the author(s) only and do not necessarily reflect those of the European Union. Neither the European Union nor the granting authority can be held responsible for them.

## Author contributions

Conceptualization: MCar, MR; Data curation: MCar, FJSL, SM, LM; Formal analysis: MCar, FJSL, AE, NR, DC; Funding acquisition: GN, MD, AB, MR; Investigation: MCar, MCas, LA, LS; Methodology: MCar, FJSL, MR; Project administration: MR; Resources: GN, MD, VVV, MRD, AB; Software: FJSL, SM; Supervision: MR; Validation: NR, VO; Visualization: MCar, FJSL, VO; Writing – original draft: MCar, MR, FJSL; Writing – review & editing: SM, MD, AB.

## Competing interests

FJSL is supported through the ENTRY-DM MSCA Doctoral Network, in which Genartis S.r.l. is a beneficiary institution. MR and MD are founders of Genartis S.r.l.. The other authors declare that no competing interests exist.

## REFERENCES

1. Kaplun, L. et al. ONT in Clinical Diagnostics of Repeat Expansion Disorders: Detection and Reporting Challenges. Int. J. Mol. Sci. 26, 2725 (2025).

2. Chen, Z. et al. Repeat expansion disorders. Pract. Neurol. 25, 204–216 (2025).

3. Rajan-Babu, I.-S., Dolzhenko, E., Eberle, M. A. & Friedman, J. M. Sequence composition changes in short tandem repeats: heterogeneity, detection, mechanisms and clinical implications. Nat. Rev. Genet. 25, 476–499 (2024).

4. Depienne, C. & Mandel, J.-L. 30 years of repeat expansion disorders: What have we learned and what are the remaining challenges? Am. J. Hum. Genet. 108, 764–785 (2021).

5. Ibañez, K. et al. Increased frequency of repeat expansion mutations across different populations. Preprint at 10.1101/2023.07.03.23292162 (2024).

6. Aston, A. N. & Dion, V. Interruptions impact clinical features of repeat expansion diseases, but how are they gained and lost? Trends Genet. TIG 41, 1056–1067 (2025).

7. Erdmann, H. et al. Repeat-associated ataxias in a German patient cohort analysed by targeted parallel long-read sequencing. Brain J. Neurol. 149, 993–1006 (2026).

8. Maestri, S. et al. Navigating triplet repeats sequencing: concepts, methodological challenges and perspective for Huntington’s disease. Nucleic Acids Res. 53, gkae1155 (2025).

9. Meola, G. Myotonic dystrophy type 2: the 2020 update. Acta Myol. Myopathies Cardiomyopathies Off. J. Mediterr. Soc. Myol. 39, 222–234 (2020).

10. Botta, A. et al. A 14-Year Italian Experience in DM2 Genetic Testing: Frequency and Distribution of Normal and Premutated CNBP Alleles. Front. Genet. 12, 668094 (2021).

11. Day, J. W. et al. Myotonic dystrophy type 2: molecular, diagnostic and clinical spectrum. Neurology 60, 657–664 (2003).

12. Leitão, E., Schröder, C. & Depienne, C. Identification and characterization of repeat expansions in neurological disorders: Methodologies, tools, and strategies. Rev. Neurol. (Paris) 180, 383–392 (2024).

13. Alfano, M. et al. Characterization of full-length CNBP expanded alleles in myotonic dystrophy type 2 patients by Cas9-mediated enrichment and nanopore sequencing. eLife 11, e80229 (2022).

14. Centofanti, F. et al. The novel (TCTG)n motif in CNBP expanded alleles: composition, dynamics and genotype-phenotype correlation in Myotonic dystrophy type 2 (DM2). Hum. Genomics 20, 87 (2026).

15. Wendlandt, M. et al. Updated Structure of CNBP Repeat Expansions in Patients With Myotonic Dystrophy Type 2 and Its Implication for Standard Diagnostics. Neurol. Genet. 11, e200220 (2025).

16. Benarroch, L. et al. Comparative Analysis of CRISPR/Cas9-targeted Nanopore Sequencing Approaches in Repeat Expansion Disorders. Genomics Proteomics Bioinformatics qzaf094 (2025) doi:10.1093/gpbjnl/qzaf094.

17. Liu, Y. et al. Repeat and haplotype aware error correction in nanopore sequencing reads with DeChat. Commun. Biol. 7, 1678 (2024).

18. Huang, Y.-T., Liu, P.-Y. & Shih, P.-W. Homopolish: a method for the removal of systematic errors in nanopore sequencing by homologous polishing. Genome Biol. 22, 95 (2021).

19. Kosugi, S. & Terao, C. Comparative evaluation of SNVs, indels, and structural variations detected with short- and long-read sequencing data. Hum. Genome Var. 11, 18 (2024).

20. Chaushevska, M. et al. Get ready for short tandem repeats analysis using long reads-the challenges and the state of the art. Front. Genet. 16, 1610026 (2025).

21. Luan, T. et al. Benchmarking short and long read polishing tools for nanopore assemblies: achieving near-perfect genomes for outbreak isolates. BMC Genomics 25, 679 (2024).

22. Eisfeldt, J., Ek, M., Nordenskjöld, M. & Lindstrand, A. Toward clinical long-read genome sequencing for rare diseases. Nat. Genet. 57, 1334–1343 (2025).

23. Thiffault, I. et al. Clinical Long-Read Sequencing Test for Genetic Disease Diagnosis. JAMA Pediatr. 179, 1355–1357 (2025).

24. Espinosa, E., Bautista, R., Larrosa, R. & Plata, O. Advancements in long-read genome sequencing technologies and algorithms. Genomics 116, 110842 (2024).

25. Lang, D. et al. Comparison of the two up-to-date sequencing technologies for genome assembly: HiFi reads of Pacific Biosciences Sequel II system and ultralong reads of Oxford Nanopore. GigaScience 9, giaa123 (2020).

26. Mizuguchi, T. et al. Complete sequencing of expanded SAMD12 repeats by long-read sequencing and Cas9-mediated enrichment. Brain J. Neurol. 144, 1103–1117 (2021).

27. Tsai, Y.-C. et al. Identification of a CCG-Enriched Expanded Allele in Patients with Myotonic Dystrophy Type 1 Using Amplification-Free Long-Read Sequencing. J. Mol. Diagn. JMD 24, 1143–1154 (2022).

28. Scholz, V. et al. Parallel in-depth analysis of repeat expansions: an updated Clin-CATS workflow for nanopore R10 flow cells. Preprint at 10.1101/2024.11.05.622099 (2024).

29. Han, Y., Jang, J.-H. & Chang, H. Targeted long-read sequencing for high-resolution repeat profiling in myotonic dystrophy type 1. Exp. Mol. Med. 58, 1203–1215 (2026).

30. DeJesus-Hernandez, M. et al. Long-read targeted sequencing uncovers clinicopathological associations for C9orf72-linked diseases. Brain J. Neurol. 144, 1082–1088 (2021).

31. Udine, E. et al. Targeted long-read sequencing to quantify methylation of the C9orf72 repeat expansion. Mol. Neurodegener. 19, 99 (2024).

32. Cumming, S. A. et al. Genetic determinants of disease severity in the myotonic dystrophy type 1 OPTIMISTIC cohort. Neurology 93, e995–e1009 (2019).

33. Gomes-Pereira, M., Bidichandani, S. I. & Monckton, D. G. Analysis of unstable triplet repeats using small-pool polymerase chain reaction. Methods Mol. Biol. 277, 61–76 (2004).

34. Aldous, S. G. et al. A CAG repeat threshold for therapeutics targeting somatic instability in Huntington’s disease. Brain J. Neurol. 147, 1784–1798 (2024).

35. Mouro Pinto, R. et al. Patterns of CAG repeat instability in the central nervous system and periphery in Huntington’s disease and in spinocerebellar ataxia type 1. Hum. Mol. Genet. 29, 2551–2567 (2020).

36. Ferguson, S. et al. Species-specific basecallers improve actual accuracy of nanopore sequencing in plants. Plant Methods 18, 137 (2022).

37. Tan, K.-T., Slevin, M. K., Meyerson, M. & Li, H. Identifying and correcting repeat-calling errors in nanopore sequencing of telomeres. Genome Biol. 23, 180 (2022).

38. Schmidt, T. T. et al. High resolution long-read telomere sequencing reveals dynamic mechanisms in aging and cancer. Nat. Commun. 15, 5149 (2024).

