## Supplementary Figure and Table for "Long-read cross-platform validation reveals novel repeat features in myotonic dystrophy type 2"

### SUPPLEMENTARY TABLE 1

| Sample ID | SMRT cell Occupancy (P1 %) | Total reads | Total HiFi reads | Total HiFi reads per sample | On *CNBP* (% of total) | Normal allele | Expanded allele |
|---|---|---|---|---|---|---|---|
| DM2-97 | 10% | 989,258 | 879,668 | 72,481 | 2,569 (3.54%) | 2,261 | 308 |
| B2 | | | | 76,370 | 3,333 (4.36%) | 2,265 | 1,068 |
| DM2-13 | | | | 97,32 | 451 (4.36%) | 377 | 74 |
| A7 | | | | 51,332 | 1,795 (3.50%) | 1,654 | 141 |
| A6 | | | | 75,004 | 2,885 (3.85%) | 2,599 | 286 |
| A5 | | | | 71,906 | 2,624 (3.65%) | 2,291 | 333 |
| A8 | | | | 63,290 | 2,381 (3.76%) | 1,988 | 393 |
| B1 | | | | 67,629 | 2,831 (4.19%) | 2,317 | 514 |

**Supplementary Table 1. Statistics of PacBio sequencing.** The table reports the SMRT Cell well occupancy during the PacBio run, the total reads and total HiFi reads produced. For each patient, it reports the number of total HiFi reads generated, the total number of "complete reads" mapping on the *CNBP* locus and fully spanning the *CNBP* repeat and, of these, those attributable to either the normal or expanded alleles. ONT reads with Q > 10 and PacBio HiFi reads were included in the analysis.

**SUPPLEMENTARY FIGURE 1**

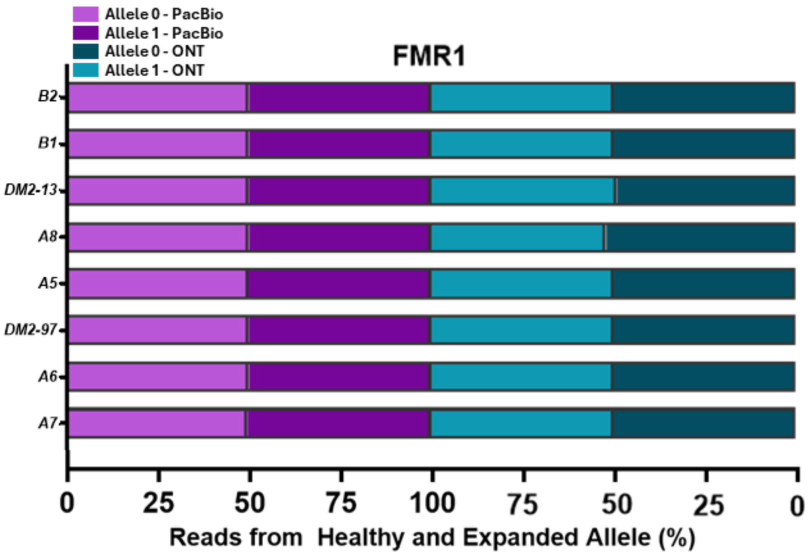


**Supplementary Figure 1. Sequencing data on the *FMR1* repeat.** Bar plots showing the proportion of reads mapping on *FMR1* and assigned to each allele using ONT and PacBio. Patients are ranked based on the median size of *CNBP* expanded allele.

**SUPPLEMENTARY FIGURE 2**

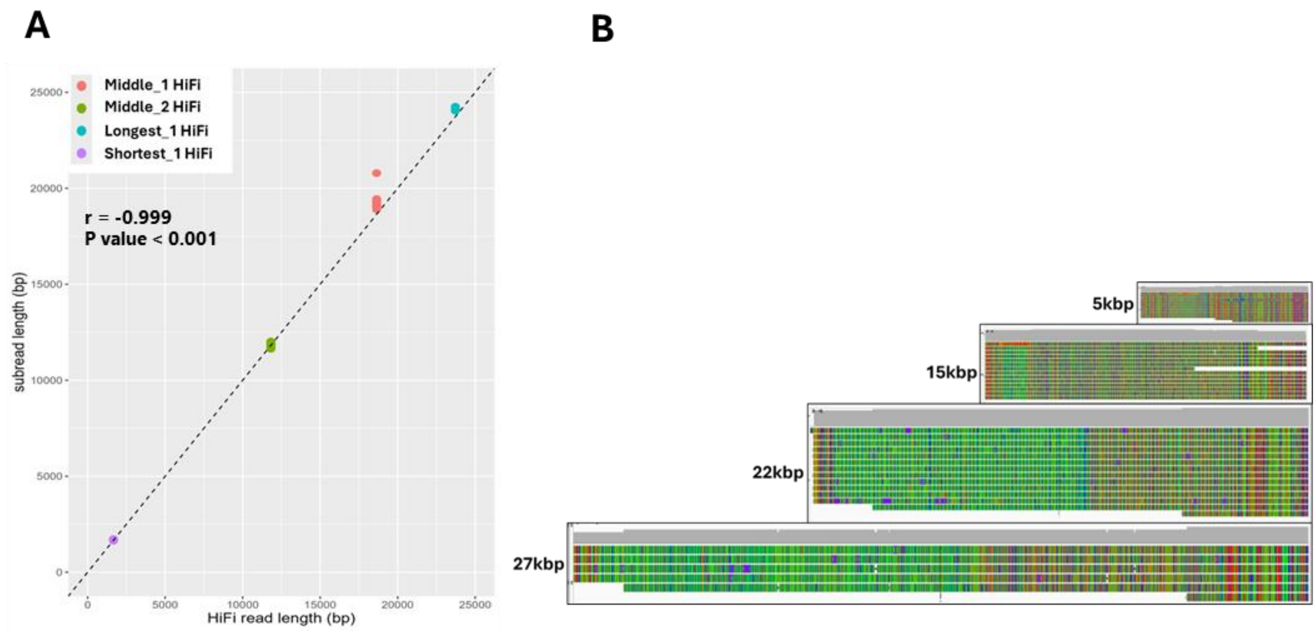


**Supplementary Figure 2. Comparison of PacBio HiFi read lengths and corresponding subreads.** (**A**) Scatter plots showing the positive and nearly-perfect correlation between the length of four representative PacBio HiFi reads from the expanded allele and the length of corresponding subreads from which they have been generated. The black dashed-line represents the bisector (x = y). Pearson correlation was used to assess the linear association between the sub-reads and HiFi reads length. (**B**) Alignments of the subreads shown in panel A on the of corresponding PacBio HiFi reads used as references.

### SUPPLEMENTARY FIGURE 3

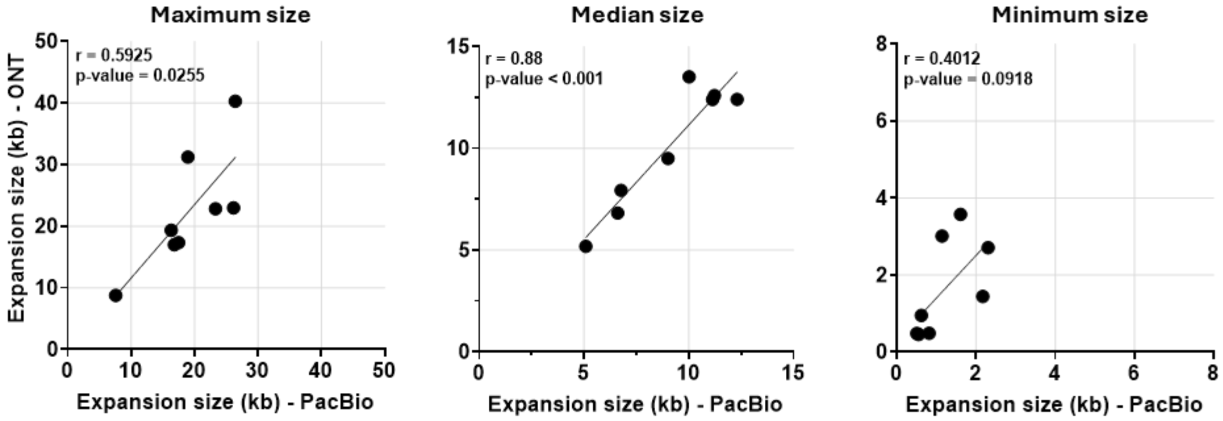


**Supplementary Figure 3. Size characterization of expanded *CNBP*-repeat alleles in DM2 patients, after down-sampling of PacBio data.** Correlations between the maximum, median, and minimum expansion lengths detected by ONT and PacBio sequencing for each DM2 patient, after downsampling PacBio data to match the number of ONT reads for each expanded allele. Pearson correlation was used to assess the linear association between the two datasets.

### SUPPLEMENTARY FIGURE 4

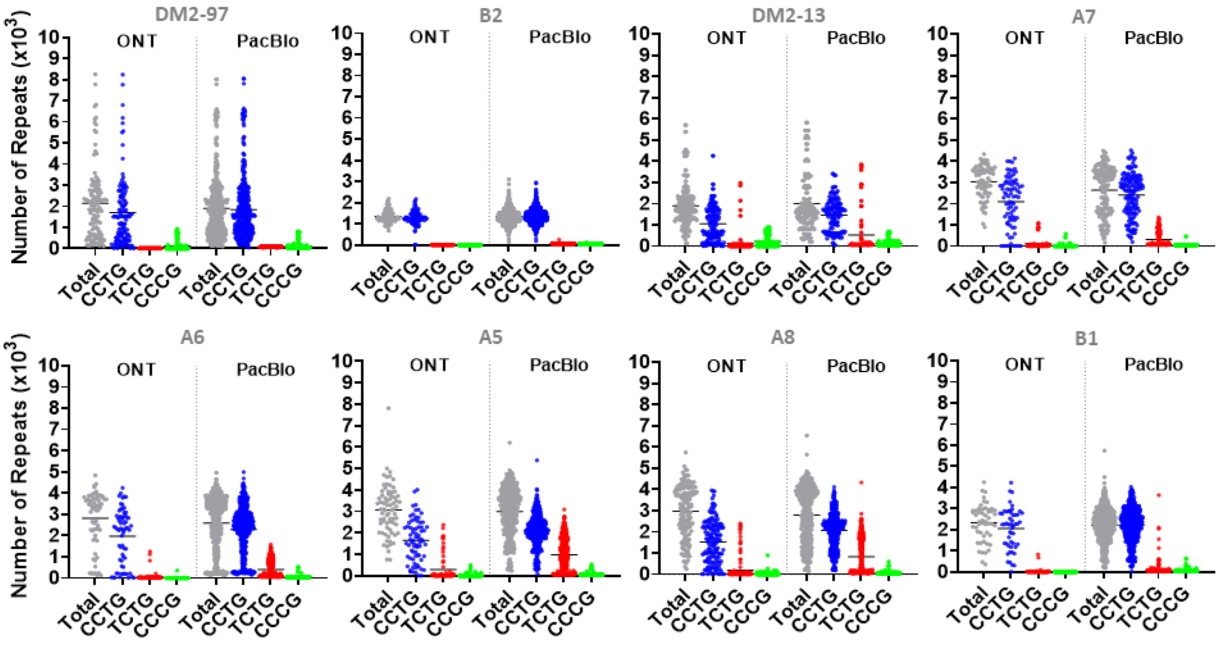


**Supplementary Figure 4. Sequence composition of expanded *CNBP* repeat alleles in DM2 patients.** The graph shows the per-read length distributions of each motif (CCTG, TCTG, CCCG) identified within reads derived from the expanded allele of each patient, based on ONT or PacBio sequencing data.

### SUPPLEMENTARY FIGURE 5

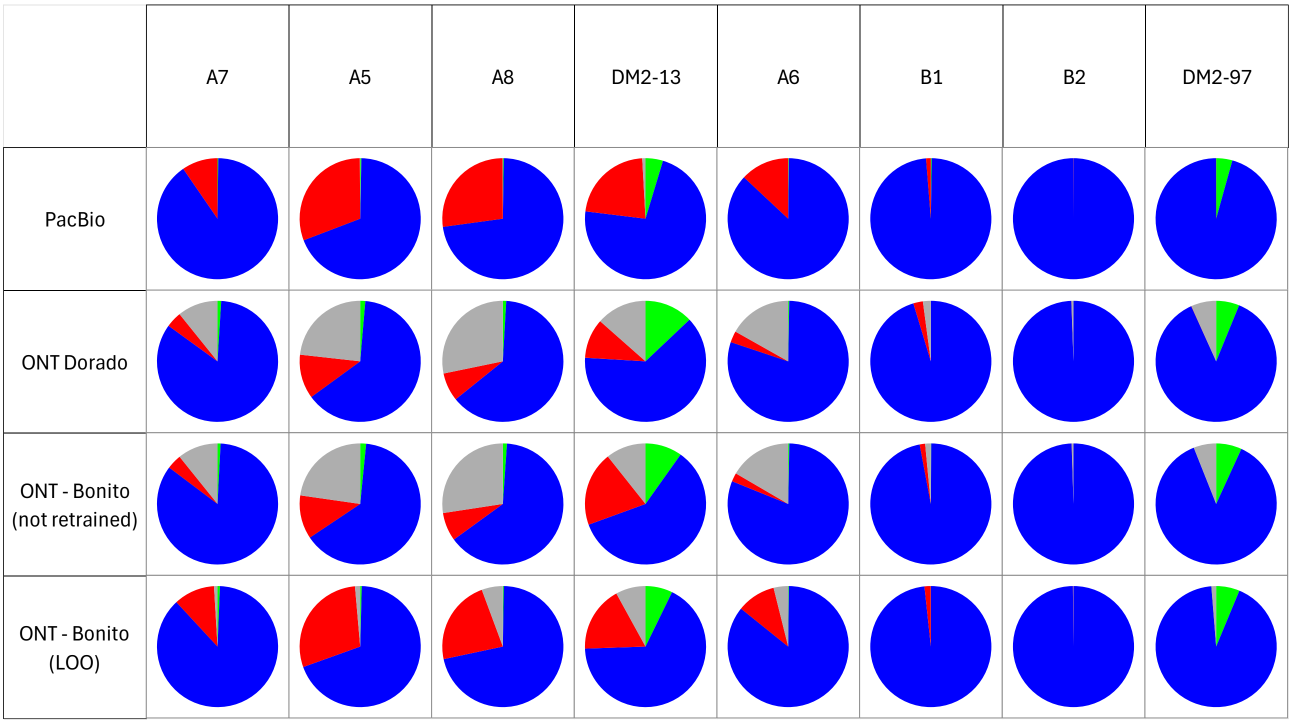


**Supplementary Figure 5. Relative abundance of *CNBP* repeat motifs in DM2 patients.** Pie charts showing the relative abundance of repeat motifs (CCTG, TCTG, CCCG) within expanded reads from each DM2 patient. The proportion of each motif is calculated as the fraction of classified repeat units (quadruplets) over the total number of repeats within reads from the expanded allele. Each horizontal line corresponds to the different sequencing platforms and basecalling approach, while columns represent an individual patient. For each patient, results are shown for PacBio data and for ONT data processed with different basecalling strategies. The Dorado and Bonito (not retrained) approaches use the same basecalling model (dna_r10.4.1_e8.2_400bps_sup@v4.3.0), while Bonito LOO used the same model retrained on PacBio data under the leave-one-out (LOO) strategy.

### SUPPLEMENTARY FIGURE 6

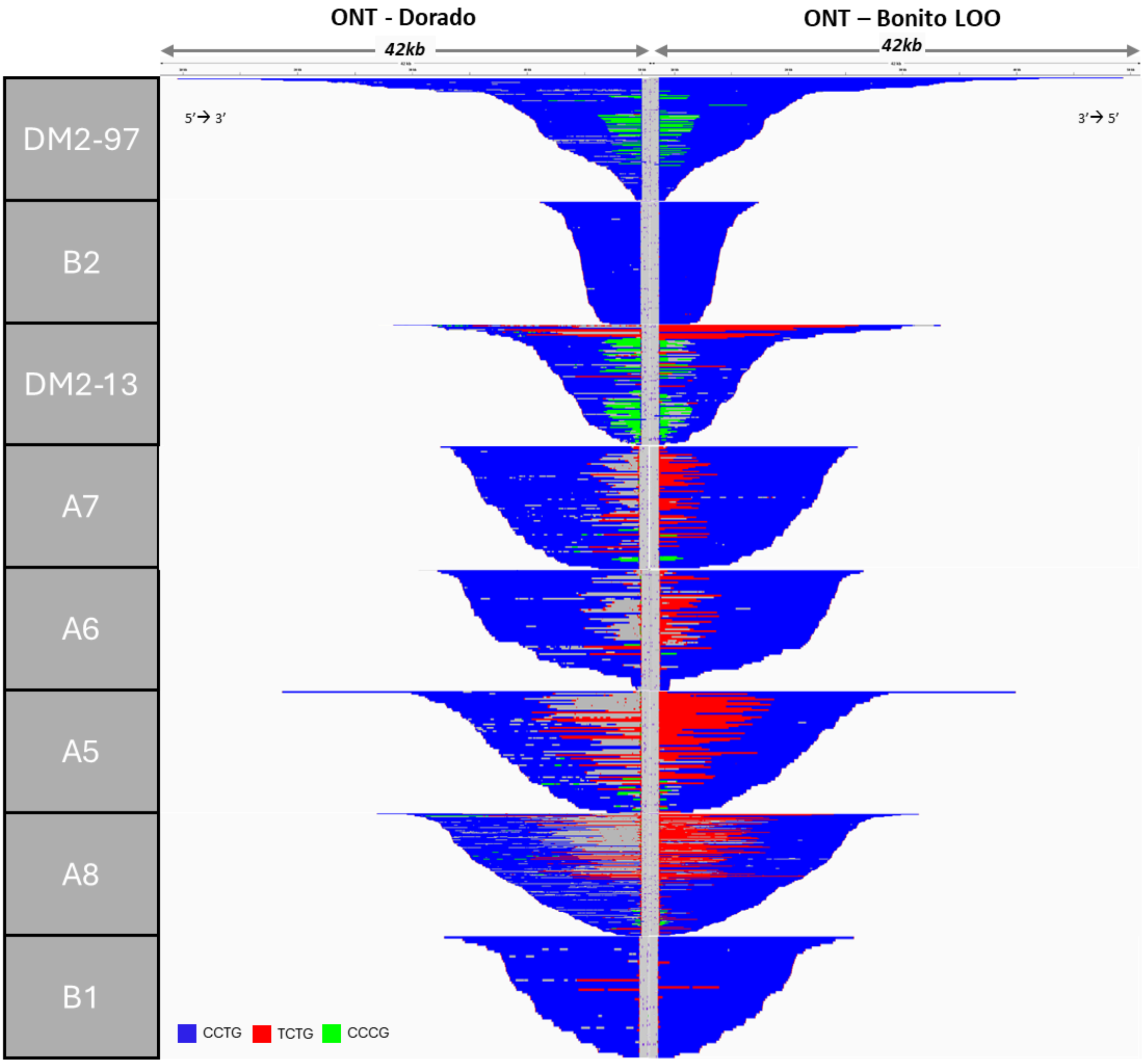


**Supplementary Figure 6. Sequence structure of expanded *CNBP* repeat alleles based on ONT data basecalled with Dorado or Bonito.** Integrative Genomics Viewer (IGV) visualization (42-kbp windows) of ONT sequencing data from expanded alleles of each DM2 patient, basecalled with Dorado (left panel) (model dna_r10.4.1_e8.2_400bps_sup@v4.3.0) or with Bonito (right panel) using the same model retrained on PacBio data under the leave-one-out (LOO) strategy. Complete reads were aligned at the 3' end of the repeat in order to identify the repeat pattern that characterizes the expanded repeat locus. Each motif in the expanded alleles was visualized using a different colour, as indicated in the key. The same data are shown in Figure 3 (ONT Dorado) and in Figure 4 (ONT Bonito LOO).

**SUPPLEMENTARY TABLE 2**

| | *FMR1* | | *DMPK* | |
|---|---|---|---|---|
| | Dorado | Bonito (LOO) | Dorado | Bonito (LOO) |
| DM2-13 | 2.59% | 1.73% | 2.74% | 1.89% |
| A7 | 1.95% | 1.36% | 2.53% | 1.58% |
| A6 | 1.93% | 1.57% | 2.15% | 1.94% |
| A5 | 2.24% | 1.60% | 2.48% | 2.33% |
| A8 | 2.33% | 1.41% | 2.97% | 1.85% |
| B2 | 1.82% | 1.44% | 2.73% | 2.08% |
| B1 | 1.95% | 1.93% | 2.28% | 2.83% |
| DM2-97 | 1.88% | 1.27% | 2.45% | 1.89% |
| **Average** | **2.08%** | **1.54%** | **2.54%** | **2.05%** |

**Supplementary Table 2. ONT error rates after basecalling with Dorado or Bonito (LOO).** For each DM2 patient, the table reports the error rate of ONT data following basecalling with Dorado (model dna_r10.4.1_e8.2_400bps_sup@v4.3.0) or with Bonito using the same model retrained on PacBio data under the leave-one-out (LOO) strategy. Error rates were calculated as the number of mismatches identified after mapping ONT reads to the hg38 reference genome.
